# Intralaboratory evaluation of luminescence based high-throughput Serum Bactericidal Assay (L-SBA) to determine bactericidal activity of human sera against *Shigella*

**DOI:** 10.1101/2020.04.03.024950

**Authors:** O. Rossi, E. Molesti, A. Saul, C. Giannelli, F. Micoli, F. Necchi

## Abstract

Despite the huge decrease in deaths caused by *Shigella* worldwide in the last decades, shigellosis is still causing over 200,000 deaths every year. No vaccine is currently available, and the morbidity of disease coupled with the rise of antimicrobial resistance renders the introduction of an effective vaccine extremely urgent. Although a clear immune correlate of protection against shigellosis has not been established yet, the demonstration of bactericidal activity of antibodies induced upon vaccination may provide one means of functionality of antibodies induced on protecting against *Shigella.* The method of choice to evaluate the complement-mediated functional activity of vaccine-induced antibodies is the Serum Bactericidal Assay (SBA).

Here we present the development and intra-laboratory characterisation of a high-throughput luminescence-based SBA (L-SBA) method, based on the detection of ATP as a proxy of surviving bacteria, to evaluate the complement-mediated killing of human sera. We demonstrated the high specificity of the assay against homologous strain without any heterologous aspecificity detected against species-related and not species-related strains. We assessed linearity, repeatability and reproducibility of L-SBA on human sera.

This work will guide the bactericidal activity assessment of clinical sera raised against *S. sonnei.* The method has the potential of being applicable with similar performances to determine bactericidal activity of any non-clinical and clinical sera that rely on complement mediated killing.

**IMPORTANCE:** *Shigella* is an important cause of diarrhoea worldwide and antimicrobial resistance is on rise, thus efforts by several groups are ongoing to produce a safe and effective vaccine against shigellosis. Although a clear immune correlate of protection has not been established, demonstration of bactericidal capacity of sera from patients immunised with *Shigella* vaccines may provide one means of protecting against shigellosis. We have developed and fully characterised a novel high-throughput L-SBA method for evaluation of functionality of antibodies raised against *S. sonnei* in human sera. This work will allow the clinical testing of human sera raised against GMMA-based and potentially all vaccines producing antibodies than can work *via* complement mediated manner.

## INTRODUCTION

Diarrheal diseases, such as shigelloses and salmonelloses, are the second leading cause of death worldwide, resulting in millions of deaths per year, mostly in developing countries (1). *Shigella* is major cause of sustained endemic bacterial diarrhoea, especially in low and middle-income countries where accessibility to clean water is restricted. Although the improvement of hygienic conditions in the last decade has dramatically reduced the burden of the disease, *Shigella* is still responsible for more than 200,000 deaths, with a third of them being in young children (1). On top of deaths in endemic countries, enteric diseases are causing diarrhoea to travellers and militaries in developed countries, further increasing the burden and the economic and social impact of them. Therefore, the huge morbidity and mortality of the disease coupled with the rise of antimicrobial resistance (2) render the introduction of a vaccine a priority for public health. Although several approaches have been tried during the years by several groups worldwide, no vaccines are licensed yet. Among the different approaches used to produce *Shigella* vaccines, many of the candidate vaccines target the serotype-specific O-Antigen (OAg) part of the lipopolysaccharide (LPS), as OAg has been identified as a key antigen recognized by the immune system after natural infection (3). In fact, although multiple immune mechanisms may provide protection against *Shigella* and are not yet fully elucidated, it is well established that antibodies directed to OAg can fix complement and kill target bacteria in a serotype-specific manner (3, 4). Genus *Shigella* is composed by four subgroups (*S. flexneri, S. sonnei, S. dysenteriae,* and *S. boydii)* and each of them, with the exception of *S. sonnei,* is composed by different serotypes, for a total of over 50 different serotypes based on the structure of the OAg, with relative prevalence of serotypes changing geographically and over time (5). As LPS antibody production can confer protection from homologous serotypes, a multivalent *Shigella* vaccine is necessary to induce antibodies to LPS OAg from multiple serotypes in order to confer broad protection.

Several approaches are currently in development to deliver the O-antigen to the immune system, including whole cell attenuated bacteria (6), vaccines in which the O-antigen are chemically-(7) or bio-conjugated to carrier proteins (8), synthetic vaccine conjugates (9), and GMMA based vaccines (10). GMMA are outer membrane exosomes from Gram-negative bacteria, genetically modified to induce hyperblebbing and to reduce the reactogenic potential of lipid A (11, 12). GMMA are easy and inexpensive to produce, and highly immunogenic (10, 13–16). The most advanced GMMA based vaccine, 1790GAHB (10) has been tested in phase I and IIa clinical trials, conducted in European (17) and endemic sites (18), and has been demonstrated to be well tolerated, immunogenic, and able to induce a strong anamnestic response after boosting (19).

On top of vaccine immunogenicity, traditionally assessed through measurement of serum antibodies *via* antigen specific ELISA, also the functionality of antibodies raised needs to be documented. Although no correlate of protection has been yet established for *Shigella,* different approaches to assess functionality of antibodies as immunological endpoint against *Shigella* have been evaluated and have been recently reviewed (20). Among them, the serum bactericidal assay (SBA) constitutes the method of choice to measure complement-mediated bacterial killing. SBA has been accepted as an *in vitro* correlate of protection for the evaluation of immunogenicity of other bacterial vaccines, such as cholera (21) and meningococcal disease (22).

The working principle of SBA relies on reconstituting *in vitro* conditions in which antibodies recognize antigen on the surface of target bacterium and bind to exogenous complement, activate the classical pathway, thus resulting in bacteriolysis and death of the target organism. The major problem with traditional SBA is that it relies on plating and counting the target bacteria. Therefore, conventional SBA has been often considered time-consuming and labor-intensive for screening large datasets and clinical samples (22). However a lot of efforts have been made in order to increase the analytical throughput of the assay, resulting in the development of both conventional (23) and non-conventional (24, 25) high throughput SBA. We have previously demonstrated the usefulness of an high-throughput SBA method based on luminescence (L-SBA) as survival readout for several pathogens (including *S. flexneri* and *S. sonnei, Salmonella* Typhimurium, Enteritidis and Paratyphi A) using both animal (24) and human sera (26). The number of viable bacterial cells surviving the complement-mediated antibody dependent killing is quantified by measuring their metabolic ATP. After bacteria lysis, ATP becomes available to trigger a luciferase-mediated reaction, resulting in a measurable luminescence signal. In L-SBA the level of luminescence detected is proportional to the number of living bacteria present in the assay wells, which is inversely proportional to the level of functional antibodies present in the serum (24). Result of the assay is the IC50, the dilution of sera able to kill half of the bacteria present in the assay, thus representing the SBA titer of the sera. We have already demonstrated the possibility to use the L-SBA to determine the bactericidal activity of sera raised against *S. sonnei* GMMA in pre-clinical models (14). Here we present the further development of this method, showing its full characterisation using human sera, and in particular sera raised against *S. sonnei* GMMA based vaccine (1790GAHB) as model. We have characterised the method intralaboratory by assessing its specificity, linearity, and precision.

The L-SBA assay described here is a useful tool for measuring functional antibodies elicited not only by GMMA based vaccines, but in general to assess *Shigella*-specific functional antibodies *in vitro,* and potentially of all vaccines that induce antibodies capable of complement mediated killing, either from preclinical and clinical sera.

## RESULTS

### Development and optimisation of L-SBA on human sera raised against *S. sonnei* GMMA

In order to determine the possibility to assess serum bactericidal activity of human sera against *S. sonnei* by L-SBA, NVGH2863, an anti-*S. sonnei* IgG human standard serum already in use to assess quantity of human antibodies raised upon vaccination with *S. sonnei* GMMA in clinical trials (17, 18), has been used to set-up the assay conditions with human sera and characterise the assay prior moving on with testing the functionality of clinical samples. Initial experiments were conducted to test the behaviour in L-SBA of NVGH2863 under experimental conditions already established with pre-clinical sera (20% exogenous baby rabbit complement (BRC) and *S. sonnei* with stabilized LPS expression *in vitro* as target bacteria). Experiments were conducted in comparison to mouse standard serum NVGH1894, already used extensively in pre-clinical studies (24), thus serving also as bridging. Assay conditions developed in pre-clinical studies resulted to be optimal also when using human sera, without detection of prozone effect when assaying sera finally diluted at 1:30 in the assay (Fig. 1A). A prozone effect is defined for a curve readout *vs* dilution (in this case luminescence vs serum dilution) a condition in which for the first points tested (the least diluted) the readout value (luminescence) is higher than readout value obtained with points highly diluted.

**Figure 1.**
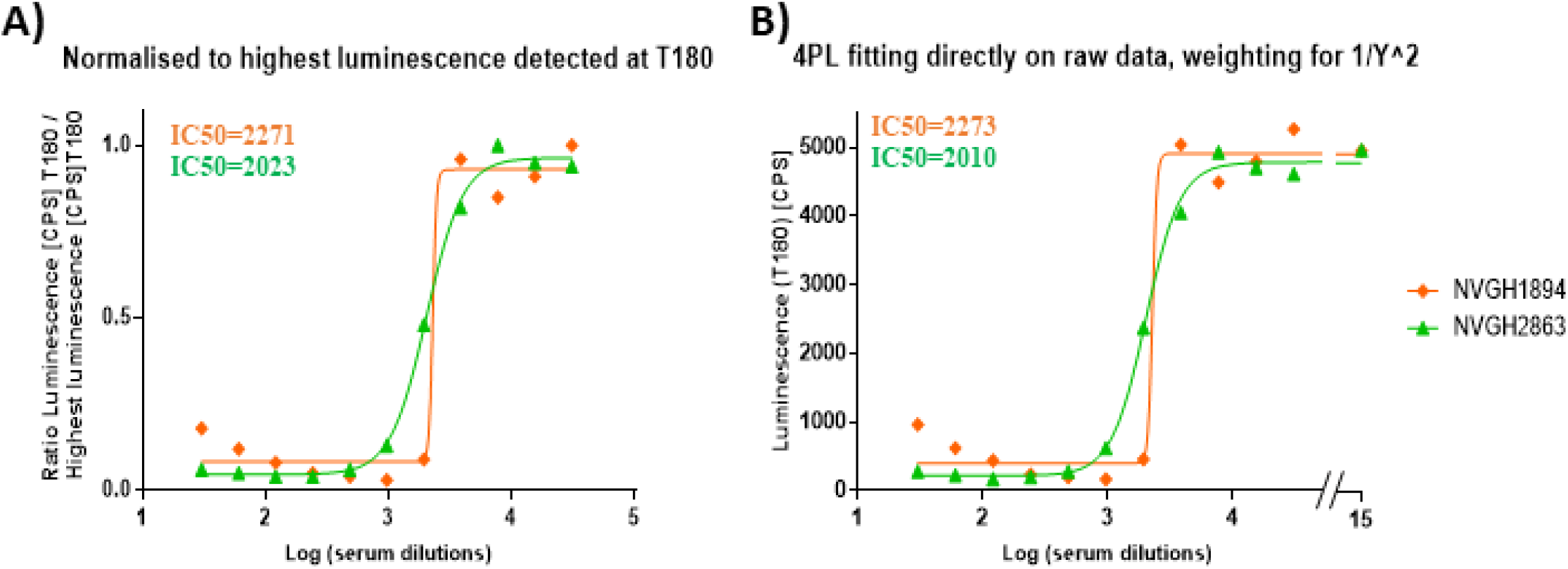
4PL fitting using different models. Representative results obtained by **A)** Fitting performed after normalisation of luminesce raw data (counts per second – CPS) for the highest luminescence detected in sera dilutions, as per (24); **B)** Fitting directly to raw luminescence (CPS), adding a weighting factor of luminescence^2 in the least mean squares calculation, assigning to luminescence detected in well with no sera an arbitrary Log dilution of 15, and forcing bottom luminescence to be between 0 and 400. Within graphs IC50 obtained testing NVGH1894 (mouse serum) and NVGH2863 (human serum) are reported in orange and green respectively.

We did an initial assessment of the homoscedasticity of the data produced at the different sera dilutions, performing 12 independent sera dilution series repeated in 6 different plates. The test for equal variance of the data obtained per each sera dilution (per each plate) confirmed the lack of homoscedasticity of the data (Fig. S1). In the 4-Parameter Logistic (PL) fitting of luminescence versus Log transformed sera dilutions, the sum of squared residuals weighted for the inverse of luminescence^2 was minimized. An improved analysis method to directly obtain SBA titres from raw luminescence data was also implemented. The aim was to minimise any raw data manipulation by operator (i.e. not to normalise for the dilution giving the highest luminescence for each dilution series, and not to individually select this value and do calculations for each sample within each run), thus reducing at minimum risk of errors. The latter is a critical aspect when testing clinical samples to ensure integrity of the data. To further improve the analysis, we also included in the 4PL fitting the luminescence value of a well with no serum, by assigning to it an arbitrary Log dilution of 15 to mimic the luminescence obtained from a serum several billions time diluted, and thus representing the maximum growth of bacteria in the assay. The use of luminescence detected on the well with no serum, coupled with mathematically forcing the 4PL regression to have a bottom luminescence below the level detected with high bactericidal sera (400 counts per second – CPS), provided solid upper and lower asymptotes of the 4PL curve fitted to data, minimising any impact of prozone (Fig. 1B).

With the identified assay conditions and improved analysis method we then moved to L-SBA characterisation by assessing precision (both in terms of repeatibility and intermediate precision), specificity, linearity, as well as to determine limit of detection and quantitation of the assay.

### Precision

Precision of the method expresses the closeness of agreement among multiple analyses of the same homogeneous representative sample tested under the prescribed conditions. It was considered at two levels: repeatability (intra-assay variation) and reproducibility as intermediate precision (inter-assay variation). ANOVA with variance component analysis (general linear model with random factors) was used to estimate the intermediate precision (defined as the variability among different days, different operators), the repeatability (defined as the variability under the same operating conditions over a short interval of time) and to evaluate the contributions of the operator and day of analysis to the variability. Thus, to assess precision of the assay, the IC50 for NVGH2863 serum was determined independently by two operators, twelve times per day, in 3 different days (72 measurements in total). Log-transformed IC50s obtained by both operators on each day have been used to determine repeatibility and intermediate precision of the assay (Fig. S2).

The analysis was characterized by an intermediate precision (CV% IP) of 6.15% and a repeatability (CV% R) of 6.15%. All the variance has been in fact attributed to the repeatibility: both day and operators resulted to be not significant (p values = 0.605 and 0.625 respectively). The average LogIC50 from all the measurements resulted to be 3.36 (or IC50 = 2528).

### Linearity

To assess linearity of the assay, NVGH2863 serum was pre-diluted in PBS (neat, 2-fold, 4-fold, 8-fold, 16-fold and 32-fold times respectively) before being probed against *S. sonnei* in L-SBA. Each pre-dilution was prepared independently by the two operators on the same day and assayed two times. We considered the average of IC50 of the undiluted serum as “true value” and from this one we calculated the expected IC50 based on the dilutons by volume performed (IC50 theoretical). A regression analysis was done on Log(IC50 experimentally obtained) vs Log(IC50 theoretical) (Fig. 2). From the analysis of variance, the linear model was significant (p < 0.001) and lack of fit not significant (p = 0.122) (Figure S3A). The residuals of the linear regression model were normally distributed (Figure S3B), the intercept was not significantly different from zero (95% CI: – 1.412; 0.096), and the slope not significantly different from 1 (95% CI: 0.856; 1.400) (Figure S3A).

**Figure 2.**
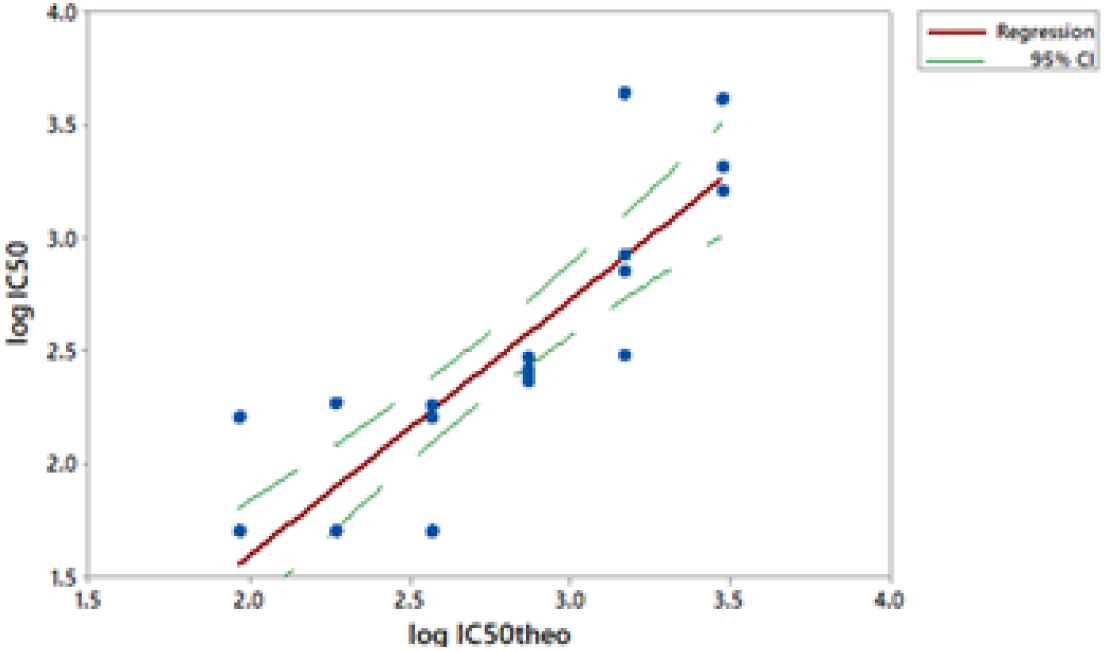
Linearity. Log(IC50 theoretical) obtained for each sample versus Log(IC50 observed) are reported. Single datapoints are indicated with blue dots. Red solid line represents the linear regression and green dashed line the 95% confidence interval (CI).

### Specificity

Specificity of the assay is the ability of an analytical procedure to determine solely the concentration of the analyte that it intends to measure. In case of *S. sonnei* 1790GAHB vaccine, the target antigen is considered the LPS OAg.

We initially assessed the homologous specificity by pre-incubating homologous *S. sonnei* purified LPS at different concentrations with test serum prior to perform the L-SBA. The aim was to determine the lowest concentration of LPS able to inhibit ≥ 70% of the IC50. Homologous competitor was spiked to NVGH2863 at the final concentrations of 50, 20, 5, 1, 0.1 μg/mL; the undepleted control was represented by NVGH2863 serum incubated with an equal volume of PBS alone. All samples were assayed in duplicate. Percentage of inhibition was determined by calculating the decrease in the observed SBA titer between samples pre-treated with competitor and undepleted control. The lowest *S. sonnei* LPS concentration, among the ones tested, that could cause a reduction of the IC50 ≥ 70% compared to the undepleted control sample resulted to be 0.1 μg/mL with over 90% depletion of the SBA titer. This concentration was then selected to assess the heterologous specificity.

A second set of experiments was performed to determine the heterologous specificity. This was carried out by pre-incubating NVGH2863 serum with an equal volume of heterologous competitor at the final concentration of 0.1 μg/mL. For heterologous specificity *S. flexneri* 1b, *S. flexneri* 2a, *S. flexneri* 3a OAg (heterologous but from the same species) and *Salmonella* Typhimurium OAg (heterologous from a different species) were tested; internal controls for these experiments were represented by serum preincubated with an equal volume of PBS alone (undepleted), and by serum preincubated with *S. sonnei* LPS (to further confirm homologous specificity). Specificity was determined as % IC50 inhibition; this was calculated using the following formula:

> (IC50 of the undepleted sample) – (IC50 of the sample pre-treated with competitor)/ (IC50 of the undepleted sample) *100.

Depletion with 0.1 μg/mL of *S. sonnei* LPS (homologous competitor) caused an inhibition of IC50 of 95%, confirming high specificity of the assay for *S. sonnei* LPS, whereas depletion with heterologous antigens resulted in an absent or a marginal (< 30%) decrease in SBA titer, suggesting the absence of any non-*S. sonnei* polysaccharide-specific response in the assay (Table 1).

**Table 1.**
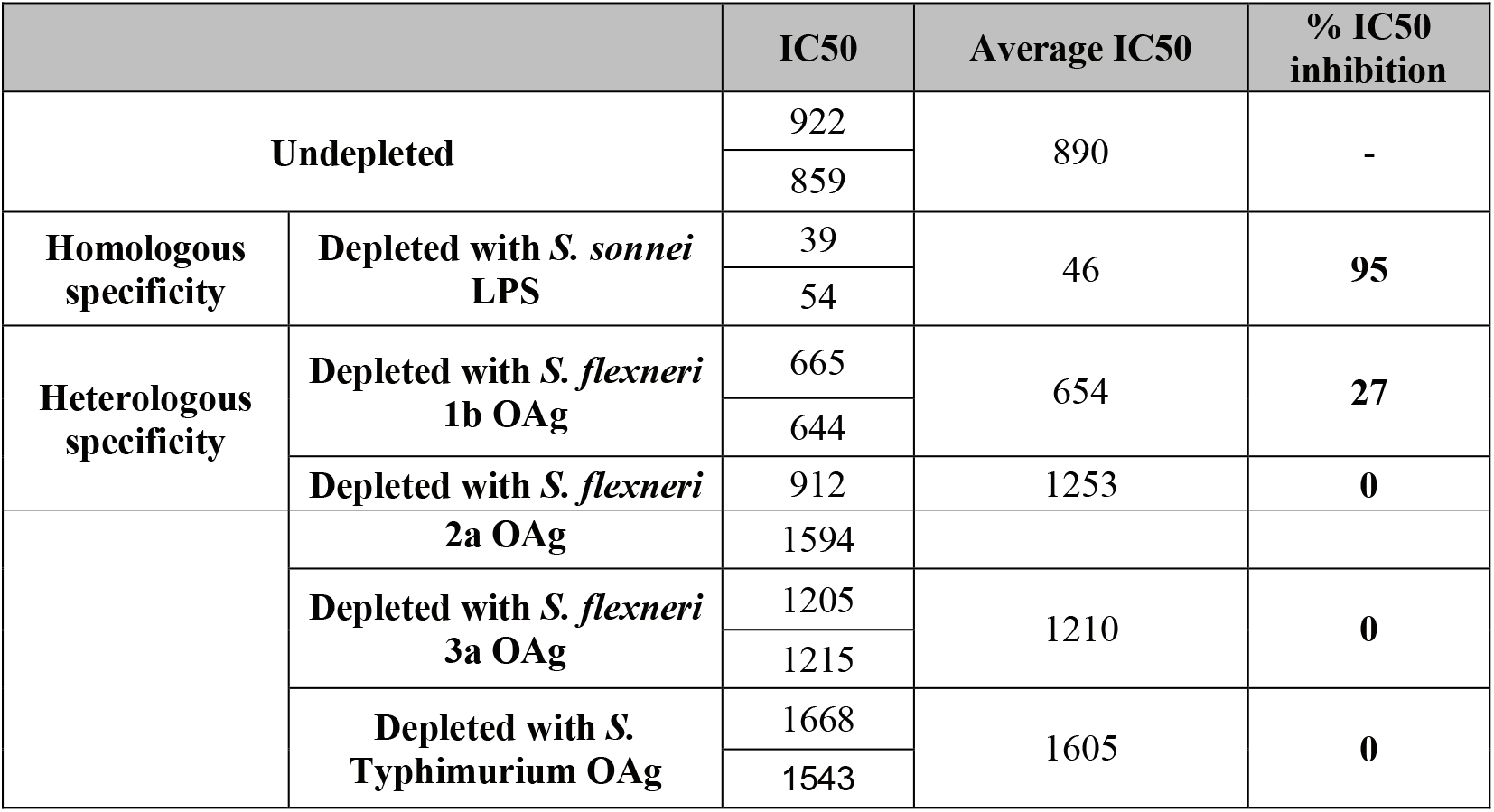
Specificity determination. Depleted samples were spiked with with 0.1 μg/mL of homologous or heterologous purified polysaccharide; undepleted samples were spiked with PBS only.

### Limit of detection (LoD) and Limit of quantitation (LoQ)

Finally, we determined the LoD and LoQ of the assay, representing respectively the lowest SBA titer than can be detected under the assay conditions, and the lowest SBA titer that can be quantified with a suitable precision. To do so NVGH2863 was pre-diluted in PBS to generate a sample with low but detectable SBA titer. These conditions simulated the worst-case scenario possible for the assay, and thus the one expected to give the highest variability. Twelve independent serial curves were tested and IC50 calculated as reported in Table 2.

**Table 2.** Determination of Limits of detection and quantitation of the assay. IC50 obtained i samples with low SBA titer.

| Repeat | 1 | 2 | 3 | 4 | 5 | 6 | 7 | 8 | 9 | 10 | 11 | 12 | Average | SD |
| --- | --- | --- | --- | --- | --- | --- | --- | --- | --- | --- | --- | --- | --- | --- |
| IC50 | 109 | 112 | 92 | 89 | 93 | 91 | 94 | 119 | 89 | 118 | 90 | 84 | 104 | 8.9 |

Limit of detection (LoD) and limit of quantitation (LoQ) of the assay were calculated accordingly to the ICH guideline Q2(R1) (27), by applying the following formulas:

> LoD = 10^(3.3 * SD)*X
>
> LoQ = 10^(10 * SD)*X

where X represents the lowest serum dilution tested in the assay (in our case = 30) and SD represents the standard deviation of IC50 obtained for the samples. LoD and LoQ resulted to be equal to an IC50 of 45 and of 100 respectively.

## DISCUSSION

The predominant readout for *Shigella* vaccine immunogenicity has been traditionally considered the serum IgG antibody level against LPS (28), that can be assessed through LPS-specific ELISA. Several assays can be considered as immunological functional readouts to determine effectiveness of antibodies raised upon vaccination, like opsonophagocytosis or serum bactericidal assay, to determine cell-mediated and cell-independent bactericidal activity of antibodies respectively (20). Although not being an established correlate of protection for *Shigella* effectiveness, ability to cause complement mediated killing has been assessed several times in sera both from convalescent patients and from vaccinated individuals (3, 4, 7). An *in vitro* assay to assess complement mediated killing represents a key indication of functional activity of antibodies raised upon vaccination with *Shigella* vaccine candidates, and this assay is traditionally represented by the SBA. The traditional SBA method used to determine the bactericidal activity of *Shigella* sera from clinical trials relies in a laborious process of plating bacteria on solid media, overnight incubation, colony counting (29), end point titer calculation without an interpolation of all sera dilutions tested (8). Such method is therefore often time consuming, highly variable, operator dependent and thus difficult to perform with consistency (Table 3). To overcome these limitations, we have recently developed an high-throughput SBA method based on luminescence readout (L-SBA), that has been already extensively described to determine the level of functional antibodies *in vitro* (24). In this study we have presented the further optimization and characterization of L-SBA on human sera. GVGH is working on developing a multivalent vaccine against *Shigella* based on GMMA technology. The most advanced *S. sonnei* component (1790GAHB), has been tested in phase I and phase II clinical trials. Immunogenicity has been evaluated so far in terms of anti-LPS IgG response induced (17, 18) (19). The work performed here will allow the analysis of the clinical samples by SBA, confirming if the antibodies elicited by 1790GAHB are able to kill *Shigella* (i.e. functionality of antibodies induced).

**Table 3.** Evaluation of troughtput between traditional and L-SBA.

|  | L-SBA | Traditional CFU-based SBA |
| --- | --- | --- |
| **Total time of execution** | 7 hours<sup>1</sup> | 1.5 working day<sup>2</sup> |
| **Data acquisition** | 2 minutes/SBA plate | 2-3 hours/SBA plate<sup>2</sup> |
| **Reproducibility** | Higher operator independency | Lower operator independency |
| **Assay throughput** | 1 operator/day: 132 individual sera in single (12 SBA plates total) | 1 operator/1.5 day: 12 individual sera in single (1 SBA plate<sup>2</sup>) |
| <sup>1</sup> to execute 1 set of 12 SBA plates<br><sup>2</sup> to execute 1 SBA plate, plating each reaction well in 1 full agar plate: 1 SBA plate corresponds to 96 agar plates |  |  |

We have successfully optimised the fitting of the data, including in data analysis a point mimicking a sera billion times diluted, allowing to establish conditions on which, without any normalisation, SBA titers can be directly obtained from raw luminescence values. The latter represents a crucial aspect to increase the throughput of the assay, but especially to reduce any potential bias due to manipulation of raw data when testing clinical samples.

In this work we have characterised L-SBA on human sera, demonstrating that in the working conditions tested, it is able to detect sera having a SBA titer as little as 45, with virtually no upper limit of detection, and to quantify with precision sera with IC50 of 100. Work is currently ongoing to further reduce LoD and LoQ. The assay showed low variability, in particular the repeatibility corresponded to the intermediate precision (CV% of 6.15%). Neither operator nor day of analysis resulted to be significant on the overall variability. Furthermore, L-SBA resulted to be highly specific for the key active ingredient of the vaccine candidate, as by depleting the serum with as little as 0.1 μg/mL of homologous LPS, over 95% reduction of IC50 was observed, whereas no depletion was observed when depleting sera with *S. flexneri* 2a, *S. flexneri* 3a and *Salmonella* Typhimurium OAg, with only a marginal SBA titer depletion (28%) observed after incubation with *S. flexneri* 1b OAg. Linearity of the assay was also assessed resulting to be good within the tested range, with a slight deflection with more diluted samples, as reported for similar assays (23). In line with that, good fitting of the data was obtained with a second order exponential model (Fig. S4).

Using the SBA configuration described here, up to 132 specimens can be tested per day by a single operator, including also a standard serum to validate each plate (12 plates can be assayed per day per operator). The analytical throughput of the described *Shigella* SBA is superior to that of another high-throughput assay recently described by Nham et al. (23), not only in terms of number of samples that can be assayed by one operator in one day, but also in terms of not having to rely on overnight incubation of plates to enable the bacteria to grow and become colonies (Table 3). As our assay uses standard reagents and requires only a luminometer to detect ATP, L-SBA can be considered simple enough to be adopted by laboratories around the world. Although inter-laboratory variability was not evaluated in our study, in case this would be observed, results could be normalised with the use of reference serum (23).

In conclusion, L-SBA applied to human sera represents an assay fully suitable to perform clinical analysis in high-throughput. Due to its specificity and versatility L-SBA can be applied to determine bactericidal activity of clinical sera raised against different *Shigella s*erotypes, helping the development of vaccines not only in single component but also when they are in multi-component formulations. Our L-SBA method can be easily extended to other pathogens, as the method has been already demonstrated to have similar performances against a broad range of pathogens using animal samples (24, 26).

## MATERIALS AND METHODS

### Bacterial strains and reagents

Working aliquots of *S. sonnei virG::cat* (30), a strain with stabilized major virulence plasmid (pSS), thus resulting in a stabilized OAg expression *in vitro* when grown in presence of antibiotic, stored frozen at −80°C in 20% glycerol stocks, were grown overnight (16-18 hours) at 37°C in Luria Bertani (LB) medium supplemented with 20 μg/mL chloramphenicol, stirring at 180 rpm. The overnight bacterial suspension was then diluted in fresh LB medium supplemented with 20 μg/mL of chloramphenicol to an optical density at 600 nm (OD_600_) of 0.05 and incubated at 37°C with 180 rpm agitation in an orbital shaker, until reaching OD_600_ of 0.22 +/− 0.02.

Baby (3-to 4-week-old) rabbit complement (BRC) was purchased from Cederlane, stored in 10 mL frozen aliquots, thawed at 4°C overnight when used. PBS was used for serum and bacteria dilutions.

LPS was extracted from *S. sonnei* by hot phenol extraction as previously reported (10), whereas OAg was extracted from *S. flexneri* 1b, 2a 3a and from *Salmonella* Typhimurium by direct acid hydrolysis as previously reported (14). All extracted polysaccharides were fully characterised in terms of sugar content, protein and nuclei acid impurities by a combination of analytical techniques, including High-Performance Liquid Size Exclusion Chromatography (31), micro BCA and absorption at 260 nm as previously reported (14).

### Serum samples

The human serum tested was an anti-human *S. sonnei* IgG standard serum (NVGH2863) that was created by pooling sera from adult subjects immunised with 1790 GAHB in non-endemic European populations (17). NVGH2863 has been already used as standard serum for *S. sonnei* LPS IgG assessment by ELISA (17–19). Frozen 50 μL working aliquots of the serum were stored at −80°C until use. In setup experiments a standard serum obtained from mice immunised with 1790GAHB (24) was also included.

All samples tested in SBA were previously Heat Inactivated (HI) at 56 °C for 30 min to remove endogenous complement activity.

Various aliquots of HI NVGH2863 serum have been used and treated as described below to determine different assay parameters.

*Samples to assess repeatibility and intermediate precision:* each sample consists on the same HI NVGH2863 serum; 12 identical samples were assayed each day by two operators and the assay was repeated in three different days by each of the two operators independently (72 samples in total, 36 per operator, 12 on each day).

*Samples to assess limit of detection and limit of quantitation:* HI NVGH2863 was diluted 20 times v:v in PBS to generate a sample with low but detectable SBA titer (expected IC50 to be around 100). Twelve identical NVGH2863 prediluted serum samples were assayed on the same day by the same operator.

*Samples to assess linearity:* HI NVGH2863 serum was assayed neat or diluted 2, 4, 8, 16, 32-fold (v:v) with PBS prior performing the assay; samples were prepared independently by two operators on the same day, with each sample assayed twice by the same operator on the same day (4 IC50 obtained for each dilution, 2 IC50 by each operator).

*Samples to assess specificity*: two sets of samples were prepared to assess homologous and heterologous specificity of the assay using HI NVGH2863 serum diluted 1:1 (v:v) in PBS alone or PBS supplemented with different quantity of homologous or heterologous purified polysaccharides. In the first experiment HI NVGH2863 serum was spiked with homologous (*S. sonnei*) purified LPS at different final concentrations (50, 20, 5, 1, 0.1 μg/mL respectively) and compared with sample spiked 1:1 with PBS alone, incubated overnight (16-18 hours) at 4°C shaking at 200 rpm in an orbital shaker prior being tested. Each spiked sample was assayed in duplicate by the same operator on the same day. The lowest concentration of LPS between the ones tested able to inhibit >70% the IC50 was then used in a second experiment to determine the heterologous specificity. In the second experiment HI NVGH2863 serum diluted 1:1 (v:v) in PBS supplemented with *S. flexneri* 1b, *S. flexneri* 2a, *S. flexneri* 3a OAg (heterologous but from the same species) or *Salmonella* Typhimurium OAg (heterologous from a different species) was prepared and assayed in comparison to sample preincubated overnight with an equal volume of PBS alone (undepleted) and a sample preincubated with *S. sonnei* LPS (to confirm homologous specificity). All samples were incubated overnight (16-18 hours) at 4°C shaking at 200 rpm in an orbital shaker prior being tested. Each spiked sample was assayed in duplicate by the same operator on the same day.

### Luminescent-SBA (L-SBA)

Serum bactericidal assay based on luminescent readout (L-SBA) was performed in 96-well round bottom sterile plates (Corning) – the SBA plate – by incubating different dilutions in PBS of HI test sera in presence of exogenous complement (BRC) and bacteria.

HI sera were serially diluted in PBS in the SBA plate (10 μL/well). The starting dilution of each serum in the assay was 1:30 (final dilution), followed by 3-fold dilution steps up to 7 dilution points, plus one control well with no sera added. Up to 12 different sera can be assayed within each SBA plate.

Log-phase cultures (OD_600_=0.22 ± 0.02) were prepared as described above and diluted to approximately 1×10^6^ Colony Forming Unit (CFU)/mL in PBS. An adequate volume of reaction mixture containing the target bacterial cells (10 μL/well) and BRC (20 μL/well) as external source of complement in PBS medium (60 μL/well) was prepared; 90 μL/well of reaction mixture were added to each well of the SBA plate containing HI serum (final reaction volume 100 μL), mixed and incubated for 3 hours at 37°C.

At the end of the incubation, the SBA plate was centrifuged at room temperature for 10 min at 4000×g. The supernatant was discarded to remove ATP derived from dead bacteria and SBA reagents. The remaining live bacterial pellets were resuspended in PBS, transferred in a white roundbottom 96-well plate (Greiner) and mixed 1:1 v:v with BacTiter-Glo Reagent (Promega). The reaction was incubated for 5 min at room temperature on an orbital shaker, and the luminescence signal measured by a luminometer (Synergy HT, Biotek).

### Calculations

The level of luminescence detected is directly proportional to the number of living bacteria present in the wells, which is inversely proportional to the level of functional antibodies present in the serum (24).

A 4-parameter non-linear regression was applied to raw luminescence (no normalisation of data was applied) obtained for all the sera dilutions tested for each serum; an arbitrary serum dilution of 10^15^ was assigned to the well containing no sera. Fitting was performed by weighting the data for the inverse of luminescence^2 and constraining the curves to have a bottom between 0 and 400 CPS. 400 CPS is the approximate value corresponding to the lowest luminescence detected at T180 for sera in all wells in which bacteria are killed (300 CPS) plus the SD of luminescence detected on those wells (100).

Results of the assay are expressed as the IC50 (the dilution of sera able to kill half of the bacteria present in the assay), represented by the reciprocal serum dilution that results in a 50% reduction of luminescence (and thus raising 50% growth inhibition). GraphPad Prism 7 software (GraphPad Software, La Jolla, CA) was used for fitting and IC50 determination.

### Statistical analysis

Statistical analyses were performed with Minitab 18, Minitab Inc as described in results section.

### Ethical statement

The human serum pool used in this study was derived from subjects enrolled in the clinical trial registered with ClinicalTrials.gov number NCT02017899. Relevant ethics and regulatory approval was obtained from respective institutional and national ethics review committees. Written informed consent was obtained before enrollment from the subjects and the trial was designed and done in accordance with Good Clinical Practice Guidelines and the Declaration of Helsinki (17).

## CONTRIBUTIONS AND ACKNOWLEDGMENT

Conceived and designed the experiments: OR, EM, AS, FMi, FN. Performed the experiments: OR; EM. Analysed the data: OR, EM, AS, CG, FMi, FN. Contributed to the writing of the manuscript: OR, EM, AS, CG, FMi, FN. This study was undertaken at the request of and sponsored by GlaxoSmithKline Biologicals SA. GSK Vaccines Institute for Global Health (Srl) (GVGH) is an affiliate of GlaxoSmithKline Biologicals SA. This work was funded in part by a grant from the Bill & Melinda Gates Foundation (OPP1133860). The funders had no role in study design, data collection and analysis, decision to publish, or preparation of the manuscript. We thank Dr Laura Bartle Martin and Dr Audino Podda for project leadership and for critically discussion about results. We would like to thank all investigators and volunteers for participation to the French study (ClinicalTrials.gov number NCT02017899).

## CONFLICT OF INTEREST

All authors were employees of the GSK Vaccines Institute for Global Health at the time in which the study was conducted. GSK Vaccines Institute for Global Health Srl is an affiliate of GlaxoSmithKline Biologicals SA. AJS possess GSK shares. This does not alter the authors’ adherence to all Journal policies on data and material sharing.

## SUPPLEMENTARY MATERIAL

**Figure S1. Test for equal variances for homoscedasticity.** Variances (x axis) from 12 replicates versus Log sera dilutions (y axis) were plotted for each of 6 different plates tested.

**Figure S2.** ANOVA with variance component analysis obtained from the 72 individual LogIC50 produced by two operators in 3 different days, each day assaying independently twelve times the same sera.

**Figure S3. A)** Regression analysis for linearity assessment. **B)** Residual plots for LogIC50.

**Figure S4. Linearity.** Log(IC50 theoretical) obtained for each sample versus Log(IC50 observed) are reported. Single datapoints are indicated with blue dots. Red solid line represents second order exponential regression and green dashed line the 95% confidence interval (CI).

